# Using a collaborative data collection method to update life-history values for snapper and grouper in Indonesia’s deep-slope demersal fishery

**DOI:** 10.1101/655571

**Authors:** Elle Wibisono, Peter Mous, Austin Humphries

**Author notes:** Corresponding author (EW).

## Abstract

The deep-slope demersal fishery that targets snapper and grouper species is an important fishery in Indonesia. Boats operate at depths between 50-500 m using drop lines and bottom long lines. There are few data, however, on the basic characteristics of the fishery which impedes accurate stock assessments and the establishment of harvest control rules. To address this gap, we developed a collaborative data collection and recording system for species and length composition of commercial catches. The Crew-Operated Data Recording System (CODRS) involves fishers who take photos of each individual fish in the catch along with a low-cost vessel tracking system. As it relies on fisher’s collaboration and willingness to share data, CODRS is comparable with a logbook system but enables verification of species identification with greater spatial resolution. We implemented this system from 2015 to 2018 and gathered data from 251 captains and 2,707 fishing trips, which yielded more than one million individual fish, or 2,680 tons. While there were over 100 species in the fishery, we found that the top five species accounted for approximately half of the total catch. We also unveiled fifteen species previously not associated with the fishery due to the fish being eaten on-board, used as bait, or sold prior to being recorded by traders. Using these data, we updated life-history parameters (length at maturity, optimum fishing length, asymptotic length, and maximum length) of the top 50 species in the fishery based on the maximum observed length; this study resulted in higher estimates for maximum length, most likely due to the high sampling size. For some species, the discrepancies between different sources were large, whereas others were not. This collaborative data collection method and findings are useful for scientists and managers interested in conducting length-based stock assessments to establish harvest control rules for data-poor fisheries.

---

[BODY]

## Introduction

In multi-species fisheries, conventional fishery-dependent data collection methods (port sampling, logbooks, and observers) are often viewed as the best way to understand the fishery. The value of these methods to inform management, however, can be limited depending on the characteristics of the fishery and thus the quality of the data [1, 2]. Applied to tropical fisheries, many of which have high species diversity, these conventional methods suffer from problems with species identification and often cannot capture data with sufficient resolution for stock assessments. For example, port sampling requires a trained enumerator to be present at the dock the moment a fishing vessel lands fish, which usually poses a logistic challenge. In many parts of the world this is a problem because the captain is under pressure to offload the boat quickly and buyers are taking fish from the catch before the enumerator has had time to record the data. Especially for longer fishing trips, it is difficult to determine the actual fishing grounds if there is no tracking system [3]. Additionally, logbooks are difficult to enforce, and captains may be uncomfortable filling in forms that fail to reflect the workflow on board the vessel. In fact, logbooks are often completed on-shore by agents who take care of the paperwork for a boat [4]. Moreover, captains use local names for fish species, which often represent a group of species and the meaning of these local names may vary between regions [5]. Other fishery-dependent methods such as observers can only be applied on larger boats that can accommodate them, are expensive, require technical expertise, and can be unsafe due to bad working conditions [6]. These challenges are often exacerbated by a low capacity of individuals to analyze the data and make it useful for management, especially in developing countries.

Understanding the factors that characterize the deep-slope demersal fishery in Indonesia are of global importance because of the wide-reaching influence of the fishery value chain [7]. To date, however, this fishery has no accurate catch or effort data, population dynamics of target species are unknown, and vessel dynamics remain elusive (i.e. size of fleet, fishing location). Even basic data on species composition are low-resolution or inaccurate. For example, Indonesian scientific publications often misidentify the most common snapper in the deepwater demersal fishery, *Lutjanus malabaricus* as *Lutjanus sanguineus*, a species from the eastern part of the Indian Ocean [e.g., 8–10]. For some of the most common species in this demersal fishery, taxonomy is still unclear and only recently have researchers concluded that the large *Etelis* species caught in Indonesian and Australian waters is probably not *Etelis carbunculus*, but a species that has not been described yet [11]. Furthermore, official catch data from the Indonesian deep-slope demersal fishery uses species categories such as the “not elsewhere included (nei)” category that clumps many different species into one group. This categorization does not allow for stock assessments or analyses of catches based on similar biological and ecological properties.

In data-poor fisheries, length-based assessment methods are a viable way to determine fishery status and set management benchmarks [12–14]. For Indonesian fisheries specifically, length-based methods are attractive because of the relative ease to gather data on species and size composition of the catch [15]. The length-based approach focuses on four important life-history parameters: length at maturity (L_mat_), optimum fishing length (L_opt_), asymptotic length (L_inf_), and maximum length (L_max_). L_max_ is the maximum length a species can attain, L_inf_ is the mean length of fish in the cohort at infinite age, and L_mat_ is the smallest length at which 50% of the individuals in that cohort is sexually mature. L_opt_ is the length class with the highest biomass in an un-fished population. Using these life-history characteristics, catch can be assessed using three primary indicators: (i) percentage of mature fish in catch (percentage of fish > L_mat_); (ii) percent of specimens with the optimum length in catch (percentage of fish at L_opt_); and (iii) percentage of ‘mega-spawners’ in catch (percentage of fish > L_opt_) [12]. These three indicators coupled with exploitation rate, and the spawning potential ratio (SPR) can be used to inform the stock status [14, 15].

Unreliable results from previous studies create a data-gap for life history parameter values for Indonesia’s deep-slope demersal fishery. To determine life history parameters, previous studies estimate L_inf_ by using age-length data to fit the Von Bertalanffy growth function [16, 17]. These studies, however, are frequently biased due to small sample sizes, gear selectivity (not all age classes are represented in the sample), or aging error [18]. Estimation of the Von Bertalanffy parameters, however, vary depending on the inputted age range [19]. Even in large sample sizes, L_inf_ estimates could be erroneous if the growth curve is not appropriate for the species and/or the gear used for sampling has narrow selectivity [18, 20]. In fished populations, fast-growing young fish and slow-growing old fish are frequently overrepresented in size-age samples, leading to an underestimation of L_inf_ [20]. An alternative approach to estimate life-history parameters is to estimate L_max_ as the largest specimen from a large sample of fish and use it to calculate other life-history parameter values based on known relationships between the parameters [13]. However, this approach has two major challenges in the Indonesian deep-slope demersal fishery context. First, obtaining length measurements of a large sample of fish is difficult with port sampling or observers, and impossible with logbooks. Second, because of problems with species identification, it is difficult to determine whether a very large specimen of a certain species is accurate without verification.

To address these challenges, we developed a collaborative data recording system for species and length composition of commercial catches that is based on photographic records of the fish in the catch, resulting in verifiable data. This system, referred to as the Crew-Operated Data Recording System (CODRS), combines simple hand-operated cameras with GPS trackers to simultaneously record catch, time, and location. Here, we report findings from CODRS, which included 1,161,659 individual length observations, allowing us to set reliable life-history parameters for the top 50 species in the fishery based on verifiable estimations of L_max_ with large sample sizes. We also compare the accuracy of CODRS against ledger receipts to see how it differs from a more traditional fishery-dependent data collection methodology.

## Methods

### Study Area

Policy and management of Indonesia’s fisheries resources is organized using zones called Fishery Management Areas (FMA). The deep-slope demersal fishery spans multiple FMAs across different water bodies in Indonesia. Thus, in 2015 we implemented our data collection system called Crew-Operated Data Collection System (CODRS) across a wide area that included Savu and Timor Seas (FMA 573), Java Sea (FMA 712), Makassar Strait (FMA 713), Banda Sea (FMA 714), Molucca Sea (FMA 715), and the Aru and Arafura Seas (FMA 718; **Fig 1**).

**Fig 1.**
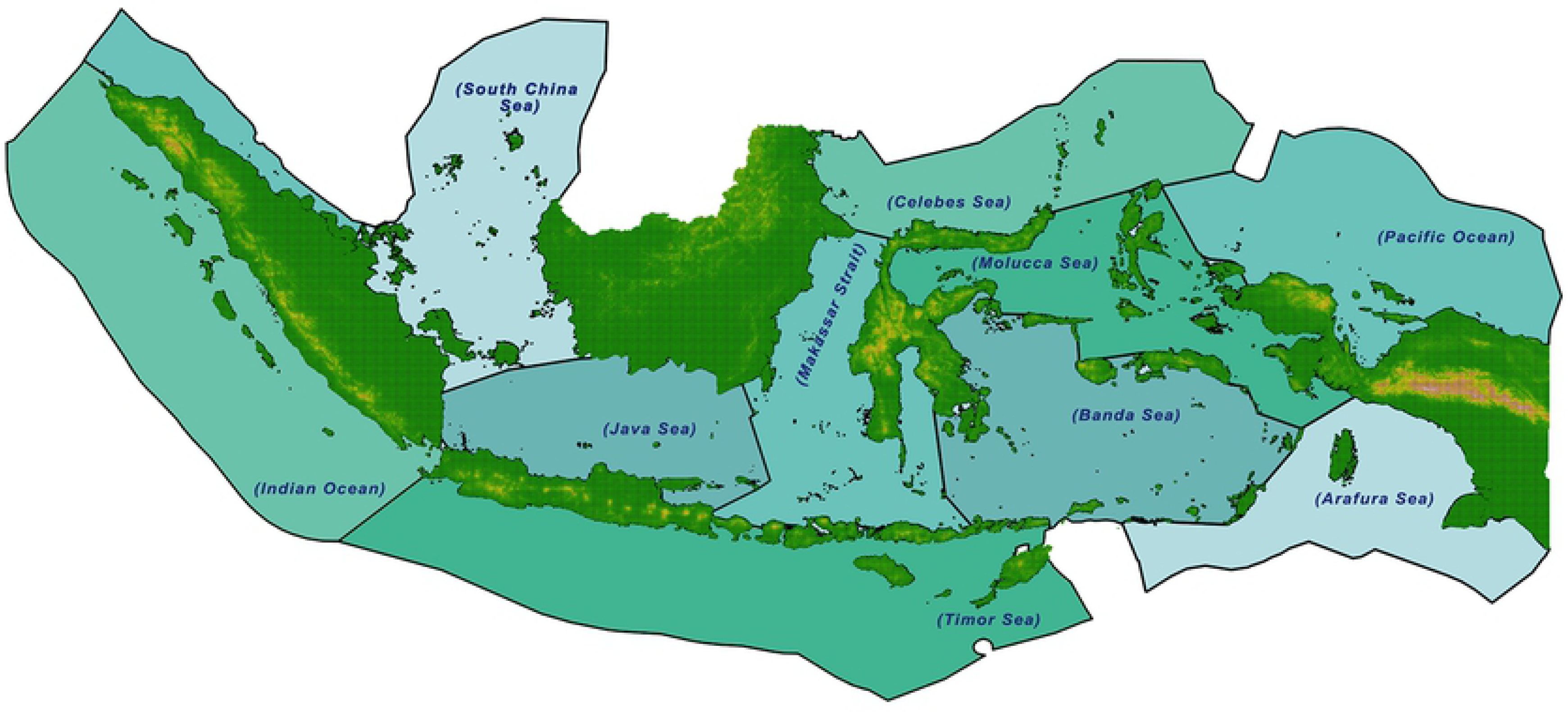
Map of the eleven Fishery Management Areas (FMA) within Indonesia. Black lines denote FMA boundaries.

Bathymetry of FMA 573, 713, 714, and 715 is characterized by mostly narrow coastal shelves, seamounts, and deep trenches. Bathymetry of FMA 712 and 718 is mostly comprised of shallow waters (50 m depth).

### Development of the Data Collection System

We recruited captains to participate in CODRS from different FMAs across the full range of the vessel sizes in the fishery (1 – 86 gross tons or GT). As an incentive for collaboration, we provided captains with monthly compensation for data collection, scaled to their vessel size category. In addition to monetary compensation, we also provided captains with a digital camera, fish measuring board, and a GPS tracking device (SPOT Trace®). We then trained captains how to take photographs of their catch and ensured the GPS tracking device transmitted the coordinates every hour. We recruited one technician per 10 vessels participating in the program (e.g., 3 technicians for 30 vessels). The technicians maintained relationships with captains and crew, and they received the digital media with the pictures from the captains after each trip. We also trained research technicians in fish identification using identification guides, frozen specimens, and photographs (**Fig. 2**).

**Fig 2.**
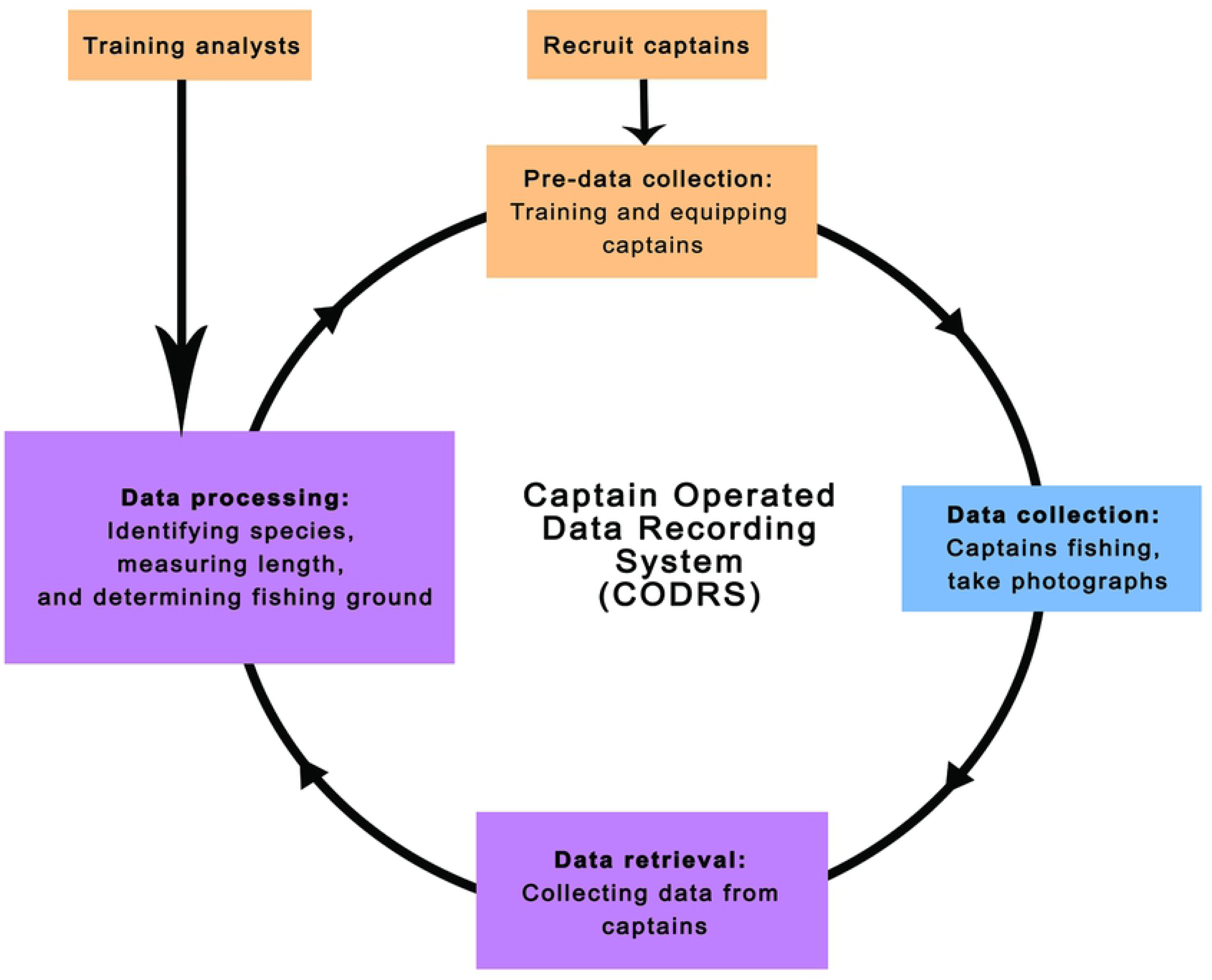
Captain Operated Data Recording System (CODRS) workflow. The system is a cycle that begins with recruitment and training of captains and analysts (orange boxes). Data is then collected at sea (blue box), then transferred to analysts for processing (purple boxes).

Data collection for each trip began when the boat left port with the GPS automatically recording vessel tracks (**Fig. 2**). After reaching the fishing grounds, crew would usually fish for a couple of hours, temporarily storing fish on the deck or in chillers. Crew would then take pictures of each fish during the packing process of putting the fish in the hold: one crew member collected fish from the deck and put it on the measuring board, where another crew member took the picture. Thereafter, the fish were stored in the hold. For very small fishing vessels (<5 GT), the process was slightly different: they took pictures upon reaching land instead of at sea. Combined with the location GPS data, the timestamps of the photographs were recorded and used to match each picture with an approximate position.

At the end of each fishing trip, which varied between two days and two months depending on vessel size, captains transferred the memory card containing the photographs of their catch to the research technicians at port. One research technician then identified fish species and another one determined the total length (TL; cm) from the pictures. An experienced third research technician examined the species identification and TL results for accuracy. A senior fisheries scientist verified the pictures of any specimens that exceed the largest fish in our database. To determine weight (kg), allometric length-weight relationships were obtained from the literature (**S1 Table**). When no values were found for a species, we used morphologically similar species to obtain the length-weight coefficients.

Catches that were abnormally low, had low quality photographs and/or only represented the first day of fishing from a multi-day fishing trip were flagged as incomplete and removed from the dataset. Catch and location data were then uploaded to a database (online) where vessel owners, captains, and researchers had access to the contents, each with different viewing privileges. For instance, captains were not able to see the fishing grounds and corresponding catches of other captains, but researchers were. Based on the quality of the photographs, research technicians provided feedback to the captains and/or crew to improve data quality on subsequent trips (**Fig. 2**).

### Data Accuracy and Catch Composition

Receipts or ledgers represented an estimate of total catch weight that was independent from CODRS. Other studies [e.g., 23] have found that sales records represent a reliable estimate of the total catch weight. To test this hypothesis, we collected receipts from fish traders that purchased fish from our partner vessels from August to November 2017. We compared these data to catch estimates from the CODRS system using paired t-tests and linear regression. Data were inspected for normality and homogeneity of variance using a Shapiro-Wilks test. We used descriptive comparisons to determine the most frequently caught species in this fishery by frequency and biomass.

### Updating Life History Parameters

Determining L_max_ values started with filtering our database for the largest fish of each species (L_x-CODRS_). Based on these values, we validated the findings by comparing L_x-CODRS_ with L_max_ documented in previous research and/or angling trophy photographs. We followed certain standards while conducting the literature review and we accepted literature values only if: (i) the study had a large sample size (n > 1000), (ii) large size range (i.e. older age classes were represented), (iii) was conducted at a comparable latitude to Indonesia, and (iv) had verifiable species identification (i.e., photograph available, species is distinct and less likely to be misidentified, species exists in the area) due to the high probability of misidentification. For studies that only estimated L_inf_ and not L_max_, we converted L_inf_ into L_max_ using the following conversion: L_max_ = L_inf_ * 1.1 [24]. Also, if fish length from literature was recorded as fork length or standard length, we converted it into total length using published conversion ratios. If L_x-CODRS_ was chosen as the new L_max_ for a species, then the photograph was reviewed by two or more research technicians and a senior fishery scientist to ensure correct species identification.

To further verify our updated L_max_ values, we searched the Internet for angling photographs for each species from comparable latitudes using key words that contained: (i) scientific name of the species of interest, (ii) scientific name of similar species, or (iii) common names from different regions. We then identified the catch species and searched for accompanying descriptive text to determine the catch area. To determine the estimated length of the fish, we used reference objects in the photograph (usually the angler’s hands) and measured the TL of the fish. Even though this approach may be less accurate, the photographs gave us a representation of the possible upper ranges of fish sizes that can help assess the plausibility of published L_inf_ or L_max_ values and the values from our CODRS database. We also compared L_mat_ values from our calculation with maturity studies that determined the length at which 50% of the population matures (of the top 15 species in the catch). We excluded studies that published values for length at first maturity. We compared L_mat_ values from areas with similar latitudes (15° S – 15° N); when not available, we included studies from other latitudes.

We calculated L_inf_, L_mat_, and L_opt_ using known relationships between the parameters and the accepted L_max_ value as described above. For all families we used L_inf_ z = 0.9 * L_max_ [22]. L_mat_ calculations differed based on the family – for Lutjanidae, L_mat_ = 0.59 * L_inf_; for Epinephelidae, L_mat_ = 0.46 * L_inf_ [23]. For other families, L_mat_ = 0.5 * L_inf_ [24]. For all families we determined L_opt_ = 1.33 * L_mat_ [25]. We then validated the results by comparing L_mat_ values with published values. We used L_mat_ estimates from histological techniques as a point of comparison because biological studies on maturation have been shown to be more robust than L_inf_ studies [26].

## Results and Discussion

### CODRS as a Method

We worked with a total of 251 captains between October 2015 and August 2018 to implement the Crew Operated Data Recording System (CODRS) in Indonesia. These captains used drop lines, bottom longlines, or a mixture of both gears. Through CODRS implementation, we obtained data from 2,707 fishing trips, which yielded 1,161,659 individual fish or 2,680 tons of catch. Vessels ranged from one to 86 GT in size. Selection of captains was roughly proportional to composition of the fleet in terms of vessel size, the Fishery Management Areas where the boat normally operates, and gear type. Because willingness of the captains to participate in the CODRS program also played an important role, catches recorded with CODRS are only roughly proportional to composition of the fleet. The dataset collected in this study includes the largest specimen ever recorded in the scientific literature and in publications on angling records for each of the 25 most common species. This is a result of the efficiency of a collaborative data collection system that involves hundreds of fishers who were able to capture verifiable data.

We used total weights from catch receipts as our control dataset to compare with CODRS. We obtained receipts from 41 captains with boats <30 GT, and from 3 captains with boats >30 GT. Because of the small sample size for large boats >30 GT, we did not use the data in our analysis. We found a statistically significant difference for the total catch weight per trip between data collected from receipts and CODRS (p < 0.001, t = 5.5243). Our CODRS dataset also recorded more fish per catch than the receipts and this became more pronounced as the catch got larger (**Fig. 3**). The estimates of total catch by CODRS appeared higher than estimates of total catch from the receipts and the variation was substantial. Receipts that indicated a total catch in the 10-500 kg range were associated with CODRS data indicating a catch of up to 1.5 metric tons. In the 500 kg - 2,500 kg per trip category, CODRS appeared to indicate a total catch that was around 50% lower than the figures indicated on the receipts. This is in contrast to the largest catches (> 2,500 kg) where there was a high correlation between CODRS and the receipts. This discrepancy was due to some fish being used as bait, eaten on-board, sold directly to individual buyers (without any receipts), or even “cheating” (rigging weighing scales to record lower weights).

**Fig 3.**
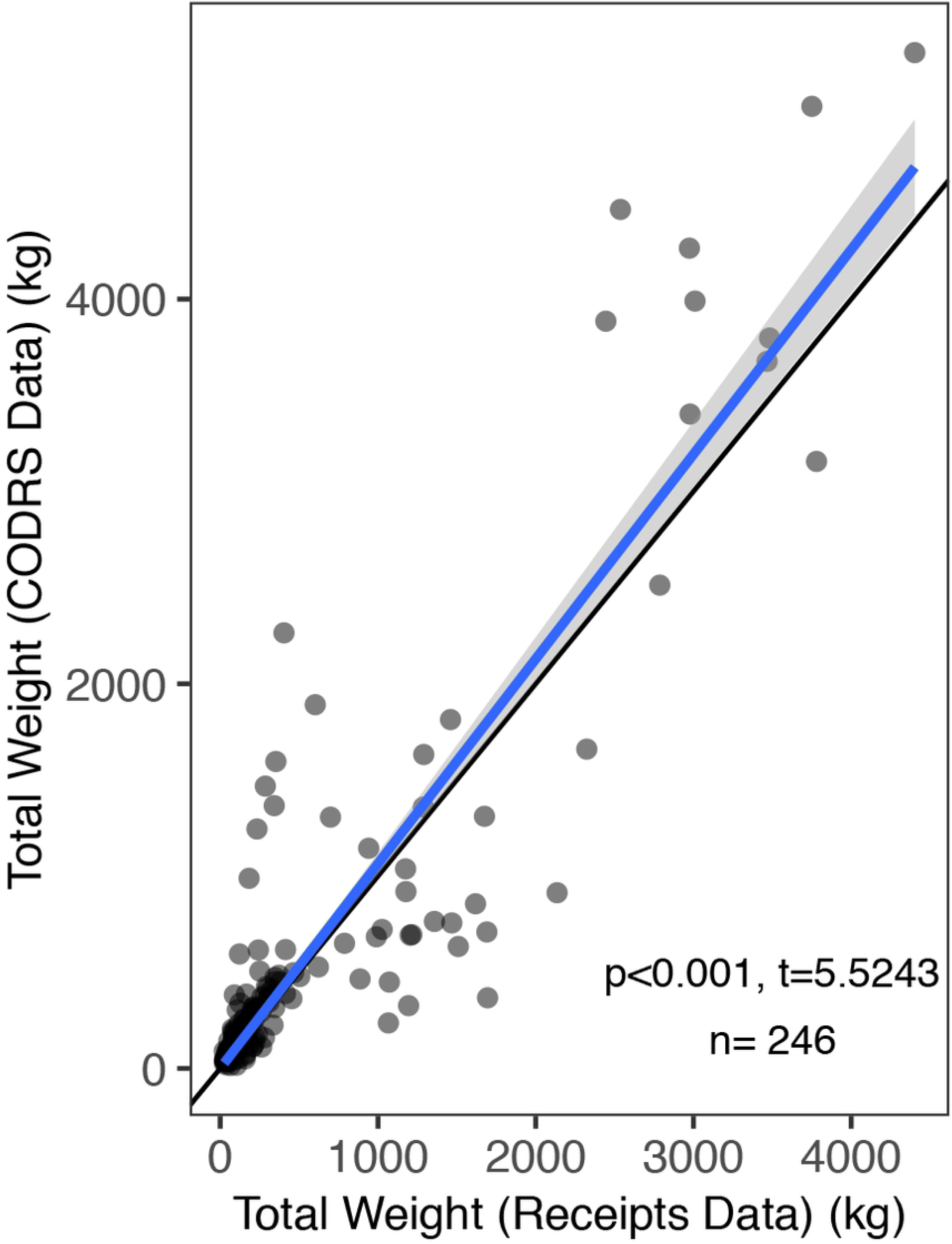
Total catch weight comparison between receipts and CODRS (Crew-Operated Data Recording System). Black line denotes 1:1 ratio between receipts and CODRS total weight; blue line denotes fitted linear regression with 95% confidence interval in grey.

It remains speculative which method provided the most accurate data for each landing, but it is remarkable that even a relatively simple observation such as total catch may easily be 20-50% higher or lower depending on the method used (ledgers versus CODRS). The problem is not with the estimation of the amount of fish in the hold at any one time. Rather, the problem is with the operational practices that affect the amount of fish in the hold as compared to the amount of fish that was actually caught. The implication is that sources of variation such as (unobserved) offloading at sea, reporting by fishers of “commercial” catch vs. catch for the local market, consumption by crew, etc., may be orders of magnitude higher than measurement errors in total catch weight at the moment that the boat is landing. These observations serve as further evidence of the importance of an on-board data collection system for this fishery as opposed to post-landing data collection methods.

The cost to implement CODRS per year was approximately $3,600-$6,300 per vessel (depending on vessel size). This is substantially more expensive than that of logbooks ($42) but not observers ($2,700 per observer trip). However, given the amount of data obtained from CODRS and its accuracy, the value of this method far exceeds that of other methods. Logbooks, observers, and CODRS all require fishers to voluntarily provide unbiased, accurate data, so this caveat is not exclusive to one method over another. One place where our CODRS method is particularly unique and useful is the detailed effort data it records for each fishing trip. Using the CODRS dataset, researchers can match GPS coordinate dates from the tracking device to the date on catch photographs, verifying time and location of catch. These parameters help to standardize catch per unit effort [27]. Researchers can also filter GPS coordinates to map fishing areas, determine the spatial distribution of fish species, analyze vessel dynamics, and determine management implications of different movement patterns [28–31].

In addition to providing catch and effort data, CODRS as a collaborative system could act as a precursor to co-management of a fishery [32, 33]. Collaborative approaches to fisheries management have gained traction in recent years as a potential solution to data-poor and open-access tropical fisheries such as those found across Indonesia [34]. This approach relies on the sharing of power and knowledge between policy-makers, researchers, and resource-users [35]. In fact, success has been shown in similar fisheries to this one which fostered collaboration and data collection for stock assessments [32, 36]. Walsh et al (2005) found that self-reporting in the Hawaii-based longline fishery for billfish can provide reliable data, provided that species identification is improved [21]. Our work on CODRS, which can be understood as a self-reporting system, corroborates this notion. In addition, CODRS resolves the species identification issue highlighted by Walsh et al. [21]. However, similar to the implementation of collaborative data collection efforts in other fisheries, communication, monitoring, and enforcement of the system is imperative to ensure data accuracy.

An advantage of CODRS over conventional data collection systems (i.e., logbooks, observers) is the ability to gather a high volume of data in a short period of time. However, despite the expedited process of data collection, the system’s success still relied on intensive data analysis, training, and monitoring captains. Thus, pre- and post-data collection efforts remain high and unavoidable given the multi-species nature of the fishery. Constant monitoring as a form of feedback is necessary to ensure compliance with the monitoring protocol [37]. In the context of the CODRS program, the most important issues that we had to address were: (i) captains needed to take photographs of their entire catch and not just a portion (including sharks and other bycatch) or their perception of the targeted catch; (ii) captains or their designated crew needed to take photographs of sufficient quality (pictures were not blurry, camera was angled properly); and (iii) captains needed to position fish on the measuring board properly. If these problems were not identified by the trained technicians, it would have led to poor data quality and misrepresentation of the catch.

We expect that technological improvements will enhance scaleability and applicability of CODRS to other fisheries. This may include things such access to cheaper high-quality cameras and an automated fish identification system [2]. Currently, photographs can be blurry especially if the photograph were taken in rough seas and/or during the nighttime. We expect that automation of image analysis through artificial intelligence will expedite the species identification process and remove may of the technical barriers to data analysis [38]. Although still in development, these technologies should soon be available and CODRS would be improved significantly, both in accuracy and cost.

### Catch Composition

Our findings show that the deep-slope demersal fishery exploited more than 100 species of fish (**S2 Table**). Half of the total catch, however, belonged to only five species (**Table 1**). The top 15 species by frequency and weight represented more than 70% of the total catch. *Lutjanus malabaricus* was the most captured species by both frequency and biomass. It contributed 19% to the total catch composition. Smaller species, such as *Epinephelus areolatus* were frequently caught, however, did not represent a large volume. Most of the catch belonged to the family Lutjanidae (snappers), subfamily Etelinae (*Pristipomoides multidens, Pristipomoides typus, Pristipomoides filamentosus, Aphareus rutilans, Etelis sp., and Prisipomoides sieboldi*). The most frequently caught species in the fishery also represented the species with highest reported economic importance [39].

**Table 1.**
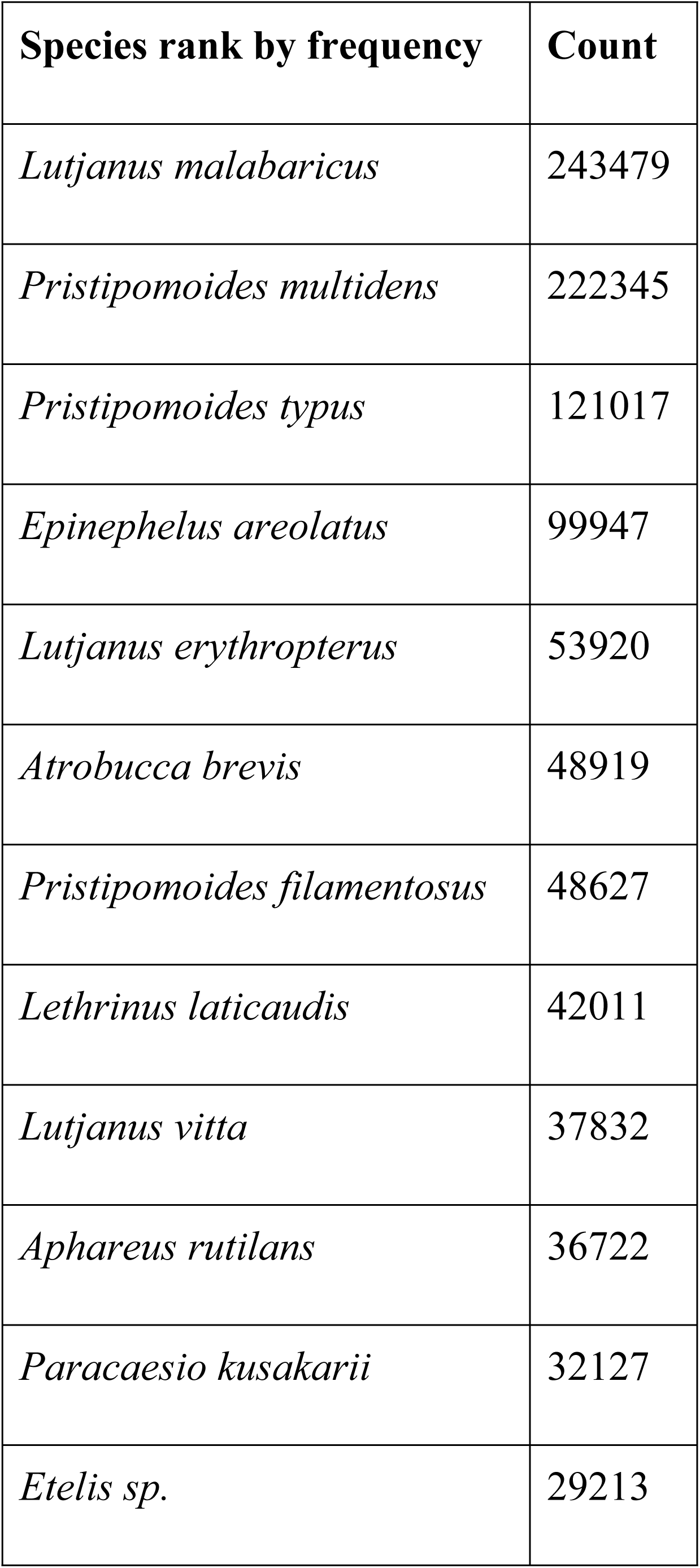
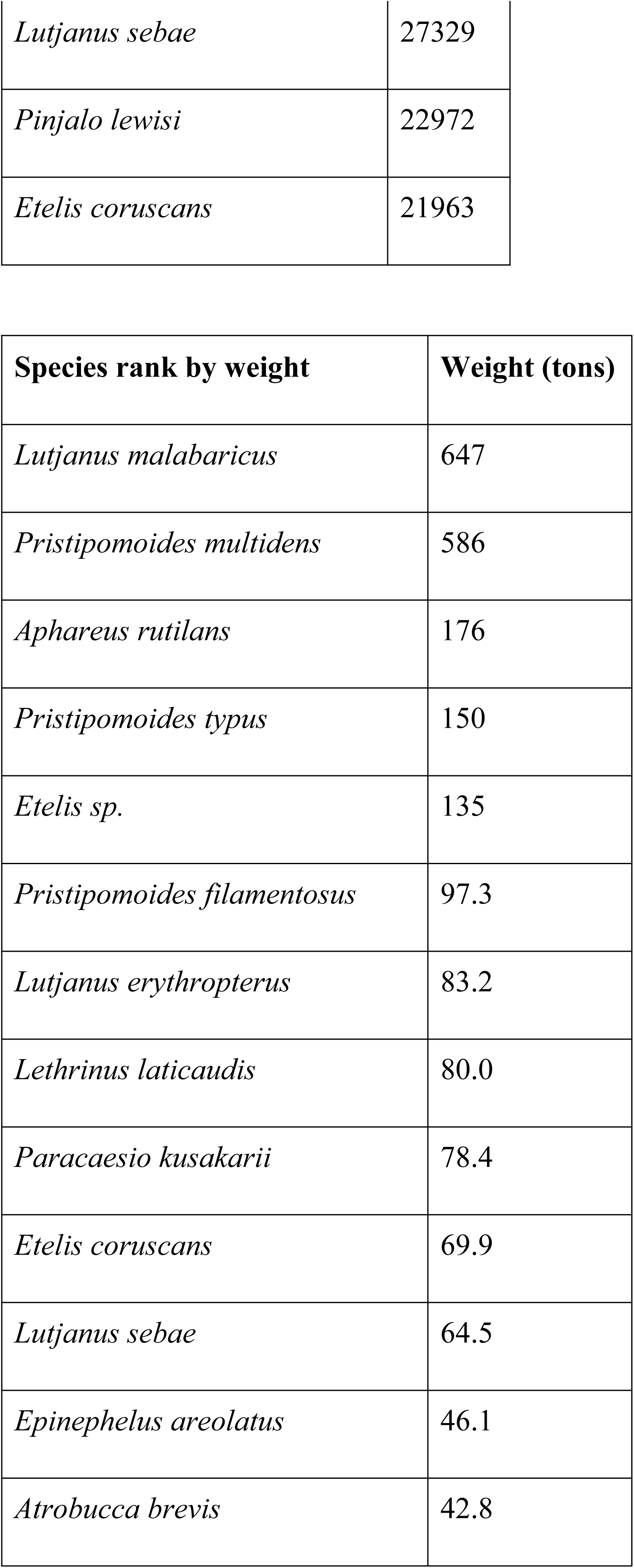
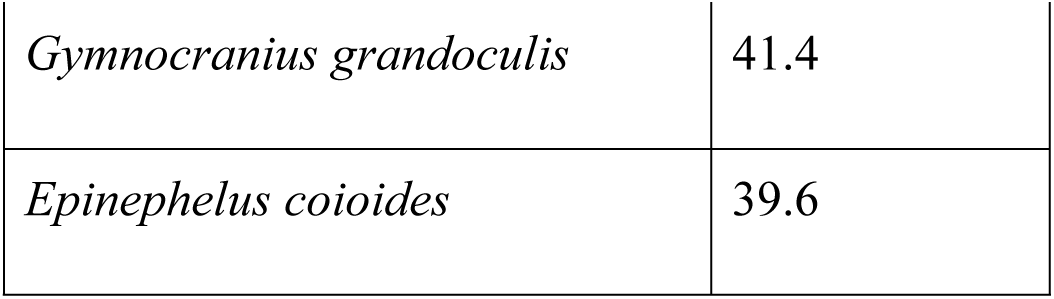
Top 15 most frequently caught species in the deep-slope demersal fishery.

Through our on-board data recording system, CODRS, we discovered 15 additional species that were not previously recorded in this fishery. These non-target catch species were either consumed, used as bait, salted on board or sold directly to the local (“wet”) market (P.Mous personal observation). This previously unreported catch consists of several species of Carangidae (*Carangoides coeruleopinnatus, Carangoides fulvoguttatus, Carangoides malabaricus, Carangoides chrysophrys, Carangoides gymnostethus, Caranx bucculentus, Caranx tille), Elagatis bipinnulata, Diagramma labiosum, Diagramma pictum, Pomadasys kaakan, Sphyraena barracuda, Sphyraena forsteri, Sphyraena putnamae,* and *Protonibea diacanthus*. In the three years of CODRS data collection, the total catch weight of these 15 species amounted to 134,470 tons. The prevalence of catches that was never offloaded and recorded on shore affirms the importance of having data collection on-board. Not only is it logistically impossible to have several enumerators or staff on shore to record catches, but the resulting data will also miss these species [5].

The dominant species in the catches of this Indonesian fishery are found throughout the deep-slope demersal fishery in the Pacific Ocean [23, 40]. However, there are differences in catch composition and properties of each species throughout the Indo-Pacific. *Etelis sp*. was recently identified as a separate species from *Etelis carbunculus* and is found throughout the Indo-Pacific but not found in Hawaii [11, 41]. In Indonesia, the ratio of *Etelis sp.* and *E. carbunculus* by count was 66 to 1 (**S2 Table**), where the average length of *Etelis sp.* in the catch was 61 cm and that of *E. carbunculus* was 41 cm. The Indonesian *Pristipomoides multidens* (the second most frequently caught species) stock does not share genetic connectivity with the adjacent Australian population [42]. *P. multidens* even has distinct genetic subdivisions within Indonesia [42]. In addition, life-history characteristics of species such as *E. carbunculus* differs between areas due to its latitudinal gradient, and ambient water temperature [41].

Different habitat and depth preferences for the major species in the catch affects species distribution in accordance with the diverse bathymetry of the area. Droplines and bottom longlines operated at different depths and habitats; dropline vessels fished at greater depths than the bottom longline. For example, *Etelis sp*., which has a depth preference between 200 to 300 m [43], were predominantly found in dropline catches. Longline vessels frequently caught non-reef species such as *Pomadasys kaakan, Diagramma pictum,* and *Diagramma labiosum,* which were rarely found in dropline catches. *Etelis sp*. and *P. filamentosus* prefer high-relief structures, such as steep drop-offs [44], and were thus captured more commonly in the dropline catches. Understanding different depth and habitat preferences of the top species in the fishery before and after maturity, along with gear-selectivity, can help inform sustainable fisheries management options such as spatial closures.

### Updating Maximum Length

Through the large number of samples and large size range per species in the CODRS database, we were able to use simple length data to derive updated L_max_ values. Our CODRS method demonstrated that it can serve as an accurate way to estimate life-history parameters by treating L_max_ and L_inf_ as biological parameters instead of a curve fitting parameter. This method was supported by robust length-frequency distributions of each species, which indicated that using L_x-CODRS_ to determine L_max_ was not an ‘anomalous’ fish; as illustrated through the length-frequency distributions of the top four species, large sizes were less prevalent, but not anomalous (**Fig 4**). Photographs of L_x-CODRS_ act as a verifiable evidence of the lengths that these species can attain. In addition, large size ranges in the database also ensured that the data collection had broad selectivity from multiple gear types and multiple vessel sizes.

**Fig. 4.**
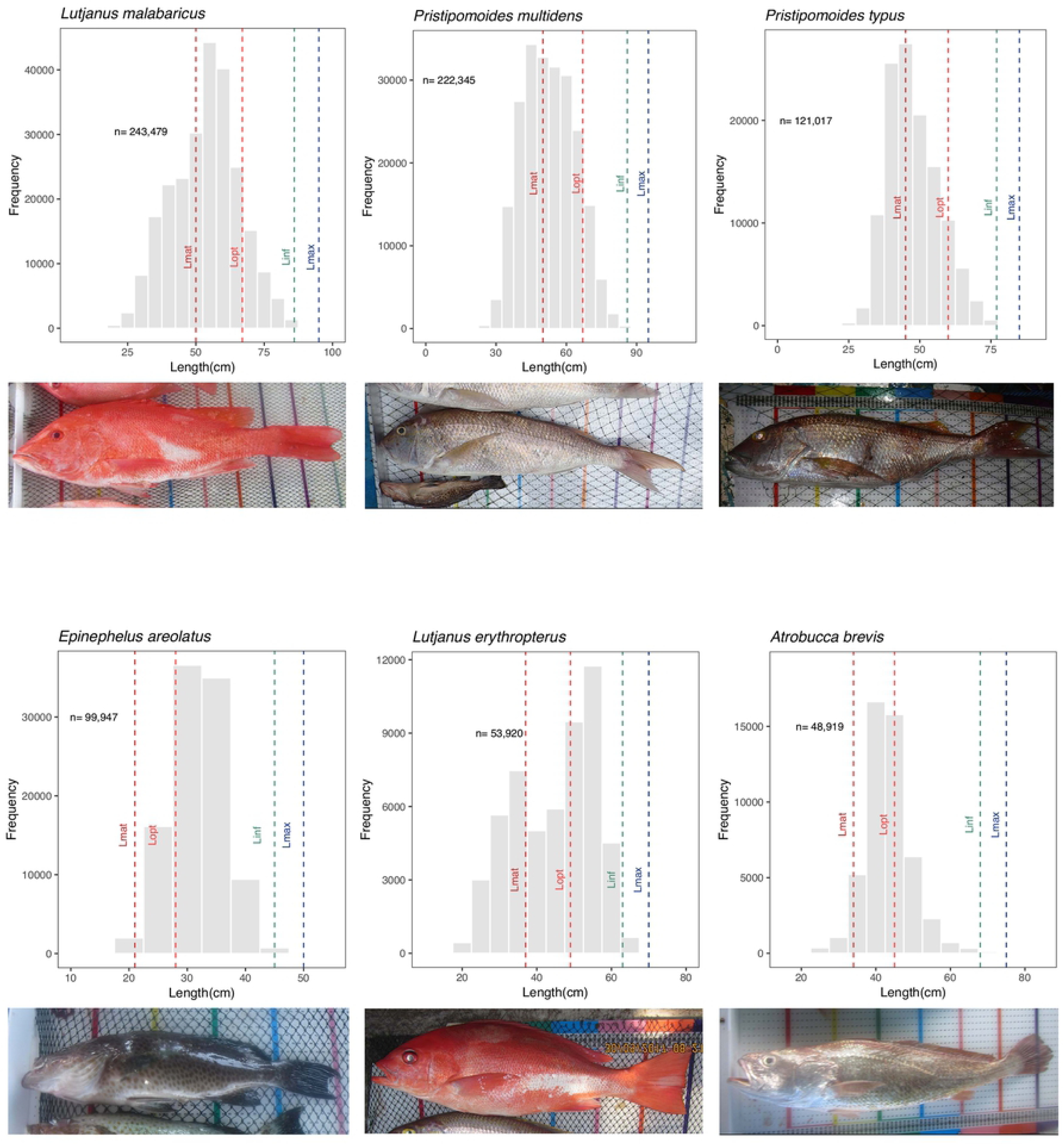
Length frequency distributions of the top six most frequently caught species in the deep-slope demersal fishery (Lutjanus malabaricus, Prisipomoides multidens, Pristipomoides typus, Epinephelus areolatus, Lutjanus erythropterus, and Atrobucca brevis). Vertical lines indicate different life history parameters. Red dashed lines represent length at maturity (L_mat_); orange dashed lines represent the length at optimum yield (L_opt_); green dashed lines represent asymptotic length (L_inf_); and the blue dashed lines represent maximum length (L_max_). Under each length-frequency distribution is a photograph from the Crew Operated Data Recording System database of the largest fish (L_x-CODRS_).

L_x-CODRS_ contributed new verifiable maximum lengths (L_max_) for the top 50 species in the fishery (**Table 2**). We did not find any L_max_ values from the literature or angling photographs that satisfied our criteria and therefore none of the updated L_max_ values are based on these data sources. Based on the L_x-CODRS_ lengths from our data, we compiled new L_max_ values that corrected for past over- or underestimation, then used this to calculate other life history parameter values (L_inf_, L_mat_, L_opt_). For some species, the discrepancies in parameter values between different sources were large, whereas others were not. For example, previous studies of *P. multidens* estimated a range of L_inf_ between 67 and 75 cm [45, 46]. As a consequence, the L_mat_ would be underestimated by 16 to 24 cm according to our data. Thus, analyzing previous research on the life-history parameters of the deep-slope demersal species required careful consideration of potential mis-identifications, or even different definitions of similar parameters. For example, some studies reported L_mat_ as the length at first maturity, whereas other studies reported L_mat_ as the length at which 50% of the population is mature [47, 48].

**Table 2.** Life history parameters (Lmat, Lopt, Linf, and Lmax) and Lx-CODRS (maximum length recorded through the Crew-Operated Data Recording System) of the top 50 most frequently caught species in the deep-slope demersal fishery.

| **Fish Species** | **L<sub>mat</sub>**<br>**(cm)** | **L<sub>opt</sub>**<br>**(cm)** | **L<sub>inf</sub>**<br>**(cm)** | **L<sub>max</sub>**<br>**(cm)** | **L<sub>x</sub>-CODRS**<br>**(cm)** |
| --- | --- | --- | --- | --- | --- |
| *Lutjanus malabaricus* | 50 | 67 | 86 | 95 | 94 |
| *Pristipomoides multidentis* | 50 | 67 | 86 | 95 | 91 |
| *Pristipomoides typus* | 45 | 60 | 77 | 85 | 85 |
| *Epinephelus areolatus* | 21 | 28 | 45 | 50 | 50 |
| *Pristipomoides filamentosus* | 48 | 64 | 81 | 90 | 88 |
| *Lethrinus laticaudis* | 29 | 39 | 59 | 65 | 63 |
| *Lutjanus erythropterus* | 37 | 49 | 63 | 70 | 70 |
| *Aphareus rutilans* | 61 | 81 | 104 | 115 | 115 |
| *Paracaesio kusakarii* | 45 | 60 | 77 | 85 | 85 |
| *Etelis sp.* | 66 | 88 | 113 | 125 | 125 |
| *Lutjanus vitta* | 24 | 32 | 41 | 45 | 43 |
| *Lutjanus sebae* | 53 | 71 | 90 | 100 | 96 |
| *Pristipomoides sieboldii* | 29 | 39 | 50 | 55 | 55 |
| *Pinjalo lewisi* | 32 | 42 | 54 | 60 | 58 |
| *Etelis coruscans* | 64 | 85 | 108 | 120 | 120 |
| *Gymnocranius grandoculis* | 36 | 48 | 72 | 80 | 76 |
| *Lutjanus timorensis* | 32 | 42 | 54 | 60 | 60 |
| *Diagramma pictum* | 38 | 51 | 77 | 85 | 81 |
| *Paracaesio stonei* | 37 | 49 | 63 | 70 | 70 |
| *Pomadasys kaakan* | 29 | 39 | 59 | 65 | 64 |
| *Wattsia mossambica* | 27 | 36 | 54 | 60 | 60 |
| *Lethrinus lentjan* | 25 | 33 | 50 | 55 | 55 |
| *Lethrinus amboinensis* | 27 | 36 | 54 | 60 | 57 |
| *Lutjanus gibbus* | 27 | 35 | 45 | 50 | 49 |
| *Protonibea diacanthus* | 61 | 81 | 122 | 135 | 130 |
| *Etelis radiosus* | 61 | 81 | 104 | 115 | 115 |
| *Carangoides chrysophrys* | 36 | 48 | 72 | 80 | 80 |
| *Lethrinus rubrioperculatus* | 20 | 27 | 41 | 45 | 45 |
| *Caranx bucculentus* | 34 | 45 | 68 | 75 | 72 |
| *Lutjanus argentimaculatus* | 50 | 67 | 86 | 95 | 95 |
| *Pinjalo pinjalo* | 42 | 56 | 72 | 80 | 77 |
| *Cephalopholis sonnerati* | 23 | 30 | 50 | 55 | 55 |
| *Caranx sexfasciatus* | 38 | 51 | 77 | 85 | 85 |
| *Paracaesio gonzalesi* | 29 | 39 | 50 | 55 | 51 |
| *Epinephelus morrhua* | 31 | 41 | 68 | 75 | 71 |
| *Erythrocles schlegelii* | 41 | 54 | 81 | 90 | 90 |
| *Aprion virescens* | 58 | 78 | 99 | 110 | 107 |
| *Epinephelus coioides* | 50 | 66 | 108 | 120 | 119 |
| *Seriola rivoliana* | 61 | 81 | 122 | 135 | 132 |
| *Lutjanus johnii* | 48 | 64 | 81 | 90 | 90 |
| *Glaucosoma buergeri* | 32 | 42 | 63 | 70 | 70 |
| *Lutjanus russelli* | 29 | 39 | 50 | 55 | 53 |
| *Lutjanus bohar* | 48 | 64 | 81 | 90 | 88 |
| *Diagramma labiosum* | 36 | 48 | 72 | 80 | 78 |
| *Lethrinus olivaceus* | 45 | 60 | 90 | 100 | 97 |
| *Paracaesio xanthura* | 29 | 39 | 50 | 55 | 52 |
| *Lutjanus bouton* | 19 | 25 | 32 | 35 | 33 |
| *Epinephelus amblycephalus* | 33 | 44 | 72 | 80 | 78 |
| *Epinephelus bleekeri* | 33 | 44 | 72 | 80 | 79 |

We found a disparity between available information in the literature and abundance of the species in the catch. For example, *P.typus*, the third most abundant species in the catch, had almost no previous studies on its life history parameters. This species is similar to and sometimes mixed with *P. multidens* during trade [49]. However, we believe that *P. typus* grows to a smaller L_max_ than *P. multidens*. The largest fish in our sample was larger than any other published values or photographs from any region. Similarly, very little literature was available on *Epinephelus areolatus* for its life history parameters or other biological characteristics despite the high recorded abundance in the catch. These disparities highlight a data gap in the literature that hampers our understanding of this lucrative and ecologically important demersal fishery.

Nadon & Ault define L_max_ as the 99th percentile of lengths in a population, apparently as a means to exclude “anomalous individuals” [22]. Whereas we agree with Nadon & Ault in their method to derive L_inf_ from an estimate of L_max_, we note that the 99th percentile of lengths depends not only on the life-history parameters of the species, but also on its exploitation status and selectivity of the fishing gear. This somewhat impairs the use of the 99th percentile of lengths as an estimate of the size that a fish can attain. Applied to the 25 most common species in the fishery, the approach of Nadon & Ault would have resulted in an estimate of L_max_ that is on average 13% lower than the largest fish we encountered (in the 25 most common species in the fishery). Upon closer inspection of the length-frequency distributions, we could not justify exclusion of the substantial range between the 99th percentile and the maximum of lengths as anomalous. We therefore adopted a more straightforward process by simply adopting the length of the largest fish encountered as the estimate for the largest size a fish can attain, from which we then derived L_inf_, accepting our estimate of L_inf_ only if it exceeded published values of L_inf_ for that species. In practice, for the 25 most common species, the L_inf_ estimates derived from our data all exceeded published L_inf_ values. We deemed 90% of L_max_ a reasonable estimate for L_inf_, acknowledging that 90% is somewhat of an arbitrary value [22].

During literature and photograph review, determining the data validity remained a challenge due to species identification issues [23]. *Aphareus rutilans* have frequently been traded as *Aphareus furca* in Indonesian fisheries (P. Mous personal observation). *A. furca* has a much smaller L_max_ and predominantly lives in shallower habitats. Only after better understanding the fishery (the fishing area, fishing depth, gear type, and distribution of the fish species) could we infer that what has been recorded as *A. furca* prior to this research was actually *A. rutilans*. Such misidentification of species obfuscates stakeholders from understanding the fishery. Description on the differences between *Etelis carbunculus* and *Etelis sp*. was fairly recent [11]. Prior to 2016, life history estimates between the two cryptic species may have originated from both species [11].

We assessed the original references for each value that is presented in FishBase during our literature search and assessment [50]. In the database, most references for L_max_ values were based on previous studies, identification guide, and angling trophy websites [50]. However, the referenced studies either did not fulfill our criteria or could not be found. Another issue was the L_max_ verification from identification guides. For example*, L. malabaricus* and *P. filamentosus* had L_max_ values in identification guides that were larger than L_x-CODRS_ [51, 52] However, due to the opacity of the number and lack of studies and/or trophy photographs to corroborate the values, we had to reject these L_max_ values from the identification guides. In addition, there were species misidentifications in the referenced angling database that were in turn referenced several times in FishBase. For example, a photograph of *P. filamentosus* was misidentified as *P. sieboldii*, leading to an abnormally large L_max_ value in FishBase.

### Updating Other Life History Parameters

Our method to calculate L_mat_ resulted in values that are within the range of published values, with a few exceptions (**Fig. 5**). When possible, we verified the validity of our updated L_mat_ values with those derived from available maturity studies, both within and outside the latitudinal range where it was caught. A common trend of L_mat_ values in the literature is the lack of consistency of values across studies, thus creating large L_mat_ ranges. For example, L_mat_ studies of *P. filamentosus* from latitudes near the equator tended to estimate larger values than values published in studies conducted in higher latitudes [53–55]. However, the opposite trend occurred in L_mat_ values for *L. sebae, L. malabaricus,* and *L. erythropterus* [16,56–62]. L_mat_ estimates from our methodology for *P. sieboldii, P. filamentosus, L. sebae, L. malabaricus, L. erythropterus*, and *Epinephelus areolatus* were somewhere in the middle of previously published ranges. Our L_mat_ estimates of *P.multidens* and *Etelis sp.* were lower than previous estimates in similar latitudes [51,56,63]. Finally, our L_mat_ estimates of *Lutjanus vitta* and *Lethrinus laticaudis* were larger than previous L_mat_ estimates. As one can see from these varied findings and comparisons across studies, there was no consistency on L_mat_ values that relate to latitudinal ranges from our study and the degree to which they agreed with other studies.

**Fig. 5.**
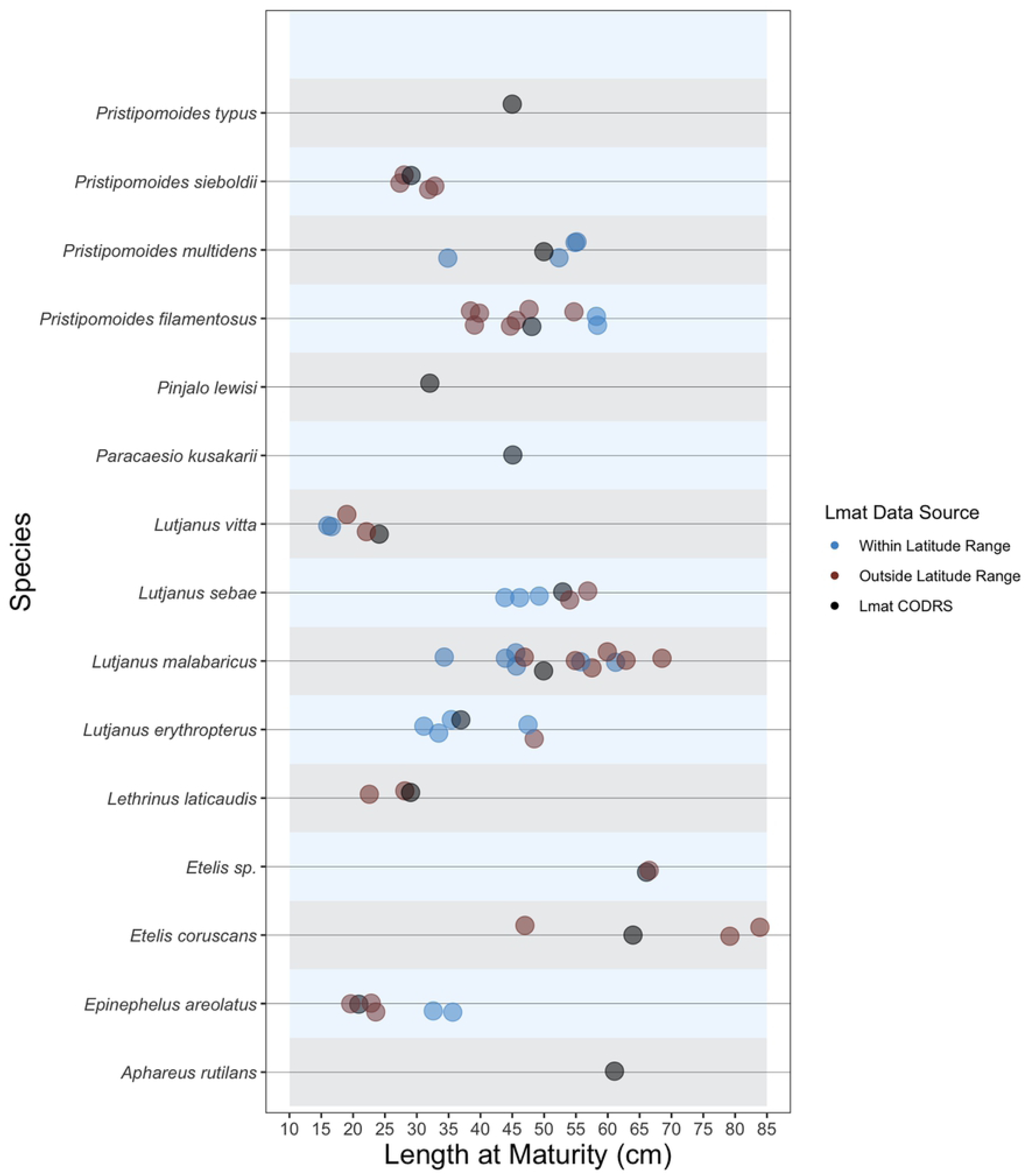
Length at maturity (L_mat_; cm) values for the top 15 species as well as Etelis sp. as calculated from our crew operated data recording system (CODRS) compared to those from the literature that were either inside or outside the latitude range of where they were caught in this study.

The differences we show between previously published L_mat_ values for the same fish species highlight the need for local values, as the difference may be important for stock assessments. For example, L_mat_ for *P*.*multidens* had the largest range of values from the literature, with 35 cm being the lowest [64] and the highest as 61 cm [56]; ours was 50 cm. Mees [53] estimated L_mat_ for *P*.*filamentosus* in Seychelles (58 cm) with samples encompassing a wide size range and large sample size. But then Ralston and Miyamoto [54] estimated L_mat_ of 44 cm from a very limited sample size. None of the previous research can represent the L_mat_ of *P.filamentosus* in Indonesian waters, however, as our estimate is in between the values proposed by the two studies. L_mat_ values for *L.laticaudis* were 22 cm (female) and 18 cm (male) [65]. These values were lower than our L_mat_ estimate, however, they originated from Shark Bay, Western Australia, which is outside the latitudinal range of our catches. The lack of previous maturity research on these species leads to high uncertainties in estimating plausible ranges for ^L^_mat_.

Similar to other life history values and studies from species-rich fisheries, species identification remains an issue. Coupled with the difficulty of acquiring samples for gonad maturity studies that is representative of the population, it is not surprising that the results of previous research were highly variable. Despite their prevalence in the catches, *P. typus, A. rutilans*, *P. lewisi*, and *Paracaesio kusakarii* did not have any maturity studies in the literature. Cross referencing values with other maturity studies were deemed important to illustrate the range of L_mat_ from pre-existing estimates and how our updated estimates compare.

### Implications for Management

Fishery-dependent data may uncover new trends in the biology of the catch that is relevant for management. In this case, the large amount of length data from CODRS helped determine new L_max_ parameters. Especially in exploited fisheries where large fish are rare, a small sample size will result in inaccurate information on the status of the stock. Life history parameter values are an integral part of length-based stock assessments. Incorrect life history parameters can lead to underestimation or overestimation of percentages of catches in each category and the status of the stock. The three indicators of overfishing (percentage of mature fish in the catch, percentage of optimum length, and percentage of mega-spawners) and other length-based stock assessment methods, such as reference points based on spawning potential ratio (SPR) and/or numerical population model rely on precise estimates of the L_mat_, L_opt_, and L_inf_ [12,13,66]. With proper interpretations, results from these assessments can inform fishery managers on the sustainability of the species in the fishery [25].

The consequence of erroneous life history parameters depends on the magnitude of the value discrepancy. Large value discrepancies may skew outcomes of stock assessments. For example, *P. typus’*s L_max_ from the FAO species catalogue was 70 cm; we estimated 85 cm. Based on our estimates, the L_mat_ should be 8 cm larger. Based on the length-frequency distribution of *P. typus* in the catch to date, we would have underestimated the percentage of immature fish by 444%, overestimated percentage of optimum length by 14%, and overestimated the percentage of mega-spawners by 74%. As a consequence, we would have concluded that the *P*. *typus* stock is in good condition – low levels of immature fish and high levels of optimum length fish in the catch. With the updated values, however, we observe a vastly different picture from the catch where 41% of the catch is immature. These results coupled with other assessment techniques will indicate the stock is being overfished. This simple example shows the importance to strive for the most accurate parameter values available that minimize biases and other stock assessment uncertainties for management.

To understand the characteristics of the catch in this fishery, examining the catch at a species level is important. However, current practices do not reflect this need – both government and private sector clump different species into arbitrary groups under a trade name or a common name in their respective databases. For example, *Lutjanus erytropterus, Pinajo pinjalo*, and *Pinjalo lewisi* are frequently grouped together as “red snapper” [49]. This grouping, without biological or ecological basis, can lead to underestimation of L_mat_ values of slower growing species. However, the L_mat_ between the largest and smallest species differs by up to 12 cm. Managing these species as one group would lead to overfishing of the largest growing species (*P. pinjalo*) and under-fishing of the smallest species (*P. lewisi*). Another example, *Paracaesio kusakarii* and *Paracaesio stonei* – differentiated morphologically only by the presence or absence of scales on the maxilla – differs 7 cm in its L_mat._ Currently they are recorded and traded under the same name, which results in growth overfishing of *P.stonei*.

## Conclusions

In Indonesia, a multi-species data collection program of this scale has never been documented before. Our crew operated data recording system (CODRS) as a method proved to be an accurate and effective system to gather catch and effort data for the deep-slope demersal fishery in Indonesia. In addition to collecting high-volume data, CODRS may also act as a first step to collaborative fishery management by engaging fishers in data collection and providing constant feedback between researcher and fisher. The quantity of verifiable length measurements enabled us to compare catch composition between gear types and update important life-history parameters such as maximum length (L_max_) and others which will be important for length-based stock assessments. We hope that the ability of CODRS to gather the high amount of species-specific catch and effort data in this pilot study can empower other fishery scientists and managers to replicate and improve this system in other data-poor multi-species fisheries.

## Acknowledgements

We would like to thank program manager (Laksmi Larasiti); all staff and field coordinators (Wawan Gede, Wahyu Dita, Geertruidha Latumeten, Helmy Wurlianty, Jersey Cumentas, Seran Abe, Nenden Siti Noviyanti, Syalomitha Hukom, Meysella Anugrah, Petronela Padja, Fatich Ubaidillah, Musa Adi, Josep Lumingas, Nandana Gozali, Dieri Tarau, Hastuti Sabang) at The Nature Conservancy Indonesia Fisheries Conservation Program for their help in data collection. We would also like to thank Dr. Jos Pet for his guidance in design and implementation of the CODRS program, and for his valuable comments on an early version of the manuscript.

## Supporting information

**S1 Table. The length-weight relationship (a and b value) and conversion factor from fork length (FL) or standard length (SL) to total length (TL) for the top species in the deep-slope snapper-grouper fishery.**

**S2 Table. The top 100 species in the deep-slope snapper-grouper fishery by total count and by total weight.**

## References

1. Chen Y, Chen L, Stergiou KI. Impacts of data quantity on fisheries stock assessment. Aquat Sci. 2003;65(1):92–8.

2. Bradley D, Merrifield M, Miller KM, Lomonico S, Wilson JR, Gleason MG. Opportunities to improve fisheries management through innovative technology and advanced data systems. Fish Fish. 2019;

3. Witt MJ, Godley BJ. A step towards seascape scale conservation: Using vessel monitoring systems (VMS) to map fishing activity. PLoS One. 2007;2(10).

4. Sondita MFA, Andamari R. A review of Indonesia’s Indian Ocean: Tuna Fisheries. ACIAR Project FIS/2001/079. 2003.

5. Yuniarta S, van Zwieten PAM, Groeneveld RA, Wisudo SH, van Ierland EC. Uncertainty in catch and effort data of small- and medium-scale tuna fisheries in Indonesia: Sources, operational causes and magnitude. Fish Res. 2017;193:173–83.

6. Mangi SC, Dolder PJ, Catchpole TL, Rodmell D, de Rozarieux N. Approaches to fully documented fisheries: Practical issues and stakeholder perceptions. Fish Fish. 2015;16(3):426–52.

7. Cawthorn DM, Mariani S. Global trade statistics lack granularity to inform traceability and management of diverse and high-value fishes. Sci Rep. 2017;7(1).

8. Ridho MR, Kaswadji RF, Jaya I, Nurhakim S. Distribusi Sumberdaya Ikan Demersal Di Perairan Laut Cina Selatan. J ilmu-ilmu Perair dan Perikan Indones. 2001;11(2):123–8.

9. Bianchi G, Badrudin M, Budihardjo S. Demersal assemblages of the Java Sea: a study bsed on the trawl surveys of the R/V Mutiara. In: Pauly D, Martosubroto P, editors. Baseline studies of biodiversity: the fish resources of Western Indonesia. ICLARM Stud.; 1996. p. 55–61.

10. Anggraeni Y. Identifikasi Dan Prevalensi Cacing Pada Saluran Pencernaan Ikan Kakap Merah (Lutjanus sanguineus) Di Pelabuhan Perikanan Nusantara Brondong Lamongan Jawa Timur. Skripsi. Universitas Airlangga; 2014.

11. Andrews KR, Williams AJ, Fernandez-Silva I, Newman SJ, Copus JM, Wakefield CB, et al. Phylogeny of deepwater snappers (Genus Etelis) reveals a cryptic species pair in the Indo-Pacific and Pleistocene invasion of the Atlantic. Mol Phylogenet Evol. 2016;100:361–71.

12. Froese R. Keep it simple: Three indicators to deal with overfishing. Vol. 5, Fish and Fisheries. 2004. p. 86–91.

13. Froese R, Binohlan C. Empirical relationships to estimate asymptotic length, length at first maturity and length at maximum yield per recruit in fishes, with a simple method to evaluate length frequency data. J Fish Biol. 2000;56(4):758–73.

14. Hordyk A, Ono K, Valencia S, Loneragan N, Prince J. A novel length-based empirical estimation method of spawning potential ratio (SPR), and tests of its performance, for small-scale, data-poor fisheries. In: ICES Journal of Marine Science. 2014. p. 217–31.

15. Prince J, Victor S, Kloulchad V, Hordyk A. Length based SPR assessment of eleven Indo-Pacific coral reef fish populations in Palau. Fish Res. 2015;171:42–58.

16. Newman SJ, Dunk IJ. Growth, age validation, mortality, and other population characteristics of the red emperor snapper, Lutjanus sebae (Cuvier, 1828), off the Kimberley coast of north-western Australia. Estuar Coast Shelf Sci. 2002;55(1):67–80.

17. MacCall AD. Quantitative Fish Dynamics. Vol. 96, Journal of the American Statistical Association. New York: Oxford University Press; 2009. 781–781 p.

18. Thorson JT, Simpfendorfer CA. Gear selectivity and sample size effects on growth curve selection in shark age and growth studies. Fish Res. 2009;98(1–3):75–84.

19. Hirschhorn K. The effects of different age ranges on estimated Bertalanffy growth parameters in three fishes and one mollusk of the northeastern Pacific Ocean. In: Bagenal T, editor. The ageing of fish. England: Unwin Bros; 1974. p. 192–9.

20. Taylor NG, Walters CJ, Martell SJD. Corrigendum: A new likelihood for simultaneously estimating von Bertalanffy growth parameters, gear selectivity, and natural and fishing mortality. Can J Fish Aquat Sci. 2011;68(8):1507–1507.

21. Walsh WA, Ito RY, Kawamoto KE, McCracken M. Analysis of logbook accuracy for blue marlin (Makaira nigricans) in the Hawaii-based longline fishery with a generalized additive model and commercial sales data. Fish Res. 2005;75(1–3):175–92.

22. Nadon MO, Ault JS. A stepwise stochastic simulation approach to estimate life history parameters for data-poor fisheries. Can J Fish Aquat Sci. 2016;73(12):1874–84.

23. Newman SJ, Williams AJ, Wakefield CB, Nicol SJ, Taylor BM, O’Malley JM. Review of the life history characteristics, ecology and fisheries for deep-water tropical demersal fish in the Indo-Pacific region. Vol. 26, Reviews in Fish Biology and Fisheries. 2016. p. 537–62.

24. Martinez-Andrade F. A Comparison of Life Histories and Ecological Aspects among Snappers (Pisces:Lutjanidae). Ph.D. Thesis. Louisiana State University, USA. Lousiana State University; 2003.

25. Cope JM, Punt AE. Length-Based Reference Points for Data-Limited Situations: Applications and Restrictions. Mar Coast Fish. 2009;1(1):169–86.

26. Brown-Peterson NJ, Wyanski DM, Saborido-Rey F, Macewicz BJ, Lowerre-Barbieri SK. A standardized terminology for describing reproductive development in fishes. Mar Coast Fish. 2011;3(1):52–70.

27. Maunder MN, Punt AE. Standardizing catch and effort data: A review of recent approaches. Fish Res. 2004;70(2-3 SPEC. ISS.):141–59.

28. Gomez C, Williams AJ, Nicol SJ, Mellin C, Loeun KL, Bradshaw CJA. Species distribution models of tropical deep-sea snappers. PLoS One. 2015;10(6).

29. Morris L, Ball D. Habitat suitability modelling of economically important fish species with commercial fisheries data. ICES J Mar Sci. 2006;63(9):1590–603.

30. Pelletier D, Ferraris J. A multivariate approach for defining fishing tactics from commercial catch and effort data. Can J Fish Aquat Sci. 2011;57(1):51–65.

31. Russo T, Parisi A, Cataudella S. New insights in interpolating fishing tracks from VMS data for different métiers. Fish Res. 2011;108(1):184–94.

32. Ticheler HJ, Kolding J, Chanda B. Participation of local fishermen in scientific fisheries data collection: A case study from the Bangweulu Swamps, Zambia. Fish Manag Ecol. 1998;5(1):81–92.

33. Prescott J, Riwu J, Stacey N, Prasetyo A. An unlikely partnership: fishers’ participation in a small-scale fishery data collection program in the Timor Sea. Rev Fish Biol Fish. 2016;26(4):679–92.

34. Nielsen JR, Degnbol P, Viswanathan KK, Ahmed M, Hara M, Abdullah NMR. Fisheries co-management-an institutional innovation? Lessons from South East Asia and Southern Africa. Mar Policy. 2004;28(2):151–60.

35. Berkes F. Evolution of co-management: Role of knowledge generation, bridging organizations and social learning. Vol. 90, Journal of Environmental Management. 2009. p. 1692–702.

36. Bell RJ, Gervelis B, Chamberlain G, Hoey J. Discard estimates from self-reported catch data: An example from the U.S. northeast shelf. North Am J Fish Manag. 2017;37(5):1130–44.

37. Gutiérrez NL, Hilborn R, Defeo O. Leadership, social capital and incentives promote successful fisheries. Nature. 2011;470(7334):386–9.

38. White DJ, Svellingen C, Strachan NJC. Automated measurement of species and length of fish by computer vision. Fish Res. 2006;80(2–3):203–10.

39. Moffitt RB, Parrish F a. Habitat and life history of juvenile Hawaiian pink snapper, Pristipomoides filamentosus. Pacific Sci. 1996;50(4):371–81.

40. Williams AJ, Loeun K, Nicol SJ, Chavance P, Ducrocq M, Harley SJ, et al. Population biology and vulnerability to fishing of deep-water Eteline snappers. J Appl Ichthyol. 2013;29(2):395–403.

41. Williams AJ, Wakefield CB, Newman SJ, Vourey E, Abascal FJ, Halafihi T, et al. Oceanic, Latitudinal, and Sex-Specific Variation in Demography of a Tropical Deepwater Snapper across the Indo-Pacific Region. Front Mar Sci. 2017;4.

42. Ovenden JR, Salini J, O’Connor S, Street R. Pronounced genetic population structure in a potentially vagile fish species (Pristipomoides multidens, Teleostei; Perciformes; Lutjanidae) from the East Indies triangle. Mol Ecol. 2004;13(7):1991–9.

43. Misa WFXE, Drazen JC, Kelley CD, Moriwake VN. Establishing species-habitat associations for 4 eteline snappers with the use of a baited stereo-video camera system. Fish Bull. 2013;111(4):293–308.

44. Oyafuso ZS, Drazen JC, Moore CH, Franklin EC. Habitat-based species distribution modelling of the Hawaiian deepwater snapper-grouper complex. Fish Res. 2017;195:19– 27.

45. Newman SJ, Dunk IJ. Age validation, growth, mortality, and additional population parameters of the goldband snapper (Pristipomoides multidens) off the Kimberley coast of northwestern Australia. Fish Bull. 2003;101(1):116–28.

46. Ralston S V, Williams HA. Depth distributions, growth and mortality of deep slope fishes from the Mariana Archipelago. Vol. NOAA-TM-NM, NOAA Technical Memorandum NMFS. Honolulu; 1988.

47. Lokani P, Pitiale H, Richards A, Tiroba G. Estimation of the unexploited biomass and maximum sustainable yield for the deep reef demersal fishes in Papua New Guinea. In: Polovina JJ, Shomura RS, editors. United States Agency for International Development and National Marine Fisheries Service Workshop on Tropical Fish Stock Assessment, 5-26 July 1989, Honolulu, Hawaii. 1990. p. 144.

48. Kikkawa BS. Maturation, spawning, and fecundity of opakapaka, pristipomoides filamentosus, in the northwestern hawaiian islands. Proc Res Inv NWHI UNIHI-SEAGRANT-MR-84-01. 1984;149–60.

49. Mous PJ, Gede W, Pet JS. Length Based Stock Assessment Of A Species Complex In Deepwater Demersal Bottom Long Line Fisheries Targeting Snappers In Indonesian Waters. Denpasar; 2018.

50. Froese R, Pauly D. FishBase [Internet]. 2012 [cited 2018 Nov 8]. Available from: http://www.fishbase.org

51. Allen GR. FAO species catalogue. Vol 6. Snappers of the world. An annotated and illustrated catalogue of lutjanid species known to date. Rome: FAO; 1985. 208 p.

52. Anderson WDJ. Lutjanidae. In: Smith MM, Heemstra PC, editors. Smiths’ Sea Fishes. Berlin: Springer-Verlag; 1986. p. 572–9.

53. Mees CC. Population biology and stock assessment of Pristipomoides filamentosus on the Mahe Plateau, Seychelles. J Fish Biol. 1993;43(5):695–708.

54. Ralston S, Miyamot GT. ANALYZING THE WIDTH OF DAILY OTOLITH INCREMENTS TO AGE THE HAWAIIAN SNAPPER, PRISTIPOMOIDES FlLAMENTOSUS Preparation of Otoliths Otolith Growth Rate and Specimen Age. Fish Bull. 1980;81(3):523–35.

55. Uehara M, Ebisawa A, Ohta I. Reproductive traits of deep-sea snappers (Lutjanidae): Implication for Okinawan bottomfish fisheries management. Reg Stud Mar Sci. 2018;17:112–26.

56. Newman SJ, Moran MJ, Lenanton RCJ. Stock assessment of the outer-shelf species in the Kimberly region of tropical Western Australia. Final report to the Fisheries Research and Development Corporation (FRDC) in Project No. 97/136. 2001.

57. McPherson GR, Squire L, O’Brien J. Reproduction of Three Dominant Lutjanus Species of the Great Barrier Reef Inter-Reef Fishery. Asian Fish Sci. 1992;5:15–24.

58. Stephenson P, Mant J. Adaptive Management of The Pilbara Trawl Fishery. 1993. 72 p.

59. Fry GC, Milton DA. Age, growth and mortality estimates for populations of red snappers lutjanus erythropterus and l. malabaricus from northern australia and eastern Indonesia. Fish Sci. 2009;75(5):1219–29.

60. Wahyuningsih P, Ernawati T. Population Parameters of Red Snapper (Lutjanus malabaricus) in Eastern Java Sea. BAWAL. 2013;5(3):175–9.

61. Tirtadanu, Wagiyo K, Sadhotomo B. Growth, yield per recruit and spawning potential ratio of red snapper (Lutjanus malabaricus Schneider, 1801) in Sinjai and adjacent waters. J Penelit Perikan Indones. 2018;24(1):1–10.

62. Lau PPF, Li LWH. Identification Guide to Fishes in the Live Seafood Trade of the Asia-Pacific Region. Hong Kong; 2000. 137 p.

63. Lloyd JA. Demography of Pristipomoides multidens in northern Australia and a comparison within the Family Lutjanidae with respect to depth [Internet]. James Cook University; 2006. Available from: http://researchonline.jcu.edu.au/31598/

64. Min TS, Senta T, Supongpan S. Fisheries biology of Pristipomoides (Family Lutjanidae) in the South China Sea and its adjacent waters. Singapore J Prim Ind. 1977;5(2):96–115.

65. Ayvazian S, Chatfield B, Keay I. The age, growth, reproductive biology and stock assessment of grass emperor, Lethrinus laticaudis in Shark Bay, Western Australia. North Beach: Department of Fisheries Research Division, Western Australia Marine Research Laboratories; 2004. 80 p.

66. Ault, JS; Bohnsack, J.A.; Meester GA. A retrospective (1979-96) multispecies assessment of fish stocks in the florida keys. Fish Bull. 1996;

